# The Bellerophon pipeline, improving de novo transcriptomes and removing chimeras

**DOI:** 10.1101/495754

**Authors:** Jesse Kerkvliet, Arthur de Fouchier, Michiel van Wijk, Astrid T. Groot

## Abstract

Transcriptome quality control is an important step in RNA-seq experiments. However, the quality of *de novo* assembled transcriptomes is difficult to assess, due to the lack of reference genome to compare the assembly to. We developed a method to assess and improve the quality of *de novo* assembled transcriptomes by focusing on the removal of chimeric sequences. These chimeric sequences can be the result of faulty assembled contigs, merging two transcripts into one. The developed method is incorporated into a pipeline, that we named Bellerophon, which is broadly applicable and easy to use. Bellerophon first uses the quality-assessment tool TransRate to indicate the quality, after which it uses a Transcripts Per Million (TPM) filter to remove lowly expressed contigs and CD-HIT-EST to remove highly identical contigs. To validate the quality of this method, we performed three benchmark experiments: 1) a computational creation of chimeras, 2) identification of chimeric contigs in a transcriptome assembly, 3) a simulated RNAseq experiment using a known reference transcriptome. Overall, the Bellerophon pipeline was able to remove between 40 to 91.9% of the chimeras in transcriptome assemblies and removed more chimeric than non-chimeric contigs. Thus, the Bellerophon sequence of filtration steps is a broadly applicable solution to improve transcriptome assemblies.

## Introduction

Ever since its first introduction in the late 2000’s (Z. Wang, Gerstein, & Snyder, 2009), RNASeq has been a useful way to determine transcriptome-wide gene expression levels. RNAseq data are sequenced cDNA reads from transcripts that can be aligned to a reference nucleotide dataset. By counting the aligned reads, gene expression levels are calculated. This technique has the advantage over other gene expression analysis methods, such as microarrays, that no *a priori* knowledge about the dataset is required, which makes single nucleotide variant analysis or novel transcript discovery possible. RNAseq is also a useful method for differential gene expression analysis in non-model organisms, for which little transcriptomic or genomic data is available. However, RNA-seq analysis requires a reference dataset to align the reads to. This dataset can be a high quality genome or a reference transcriptome.

There are two ways to assemble a reference transcriptome. The first method is reference-based, which is done by performing an alignment of the cDNA reads to a reference genome of high quality. The assembly can be done quickly, using reasonable computational power and the transcriptome will be of high quality as long as the genome is of high quality. For transcriptomes of organisms without a reference genome, there is the second method: a *de novo* transcriptome assembly for which no reference data are required. The most commonly used *de novo* transcriptome assembler is Trinity (Haas et al., 2013). This tool uses *de Bruijn* graphs to construct contigs from overlapping cDNA reads (Grabherr et al., 2011). However, a *de novo* assembly requires high computational power and its quality is difficult to assess, because of the lack of reference DNA or RNA data to compare it to (Li et al., 2014). Sequencing errors can greatly alter the assembled transcriptome, which thus induces errors in the differential gene expression analysis (Marchant et al., 2016; Martin & Wang, 2011).

Different types of errors can occur during a *de novo* transcriptome assembly process (see Smith-Unna *et al.*, 2016, 1). For example, assembled transcripts can be incomplete, one transcript can be assembled into multiple contigs, or multiple transcripts can be fused into one contig. Chimeric sequences can occur naturally in transcriptomes (*i.e*. not the result of assembly errors), but these sequences are rare (Frenkel-Morgenstern et al., 2012). False chimeric contigs are the product of a misassembly of multiple different transcripts that have erroneously been assembled together into one contig. This can occur when *de Bruijn* graph extension is difficult due repeated regions or when two sequences are almost identical (Lima et al., 2017). There are two defined types of false chimeric sequences: 1) the contig can be composed from different isoforms of the reference transcript, which is called a self-chimera, 2) the contig can be composed from two different transcripts, which is called a multi-chimera (Yang & Smith, 2013).

Assembly errors might be identified and filtered out by mapping the cDNA reads to the assembled contigs (Smith-Unna et al., 2016). Different patterns of read coverage can be evidence for different types of errors. For example, high variation between the number of reads mapped to a contig, or the lack of reads mapping to a contig, can indicate inappropriately assembled transcripts. In general, contigs should be evenly expressed, because different parts of a correctly assembled transcript should not be differentially expressed. An uneven expression pattern is typical of false chimeric contigs. The exception is when multiple splicing variants of a gene are present in the transcriptome and assembled by the assembler.

Only a short list of tools are available to assess the quality of a *de novo* transcriptome. The tools KisSplice (Sacomoto et al., 2012), DRAP (Cabau et al., 2017), RSEM-EVAL (Li et al., 2014) and TransRate (Smith-Unna et al., 2016) all assess the quality of a transcriptome. DRAP and KisSplice are transcriptome assemblers on their own, focusing on transcriptome quality assessment and chimera removal, while RSEM-EVAL and TransRate are post-assembly tools. When working with already assembled transcriptomes that need to be optimized, RSEM-EVAL and TransRate would be a better choice, as *de novo* assembly remains a computationally intensive task that is not easily redone. RSEM-EVAL requires a reference set of transcripts, which can be from a closely related species, and uses the reference set to estimate transcript length distribution. This thus makes RSEM-EVAL not truly reference-free. TransRate is truly reference-free and only requires the sequencing reads and the assembled transcriptome.

As gene expression levels in RNA-seq experiments are determined by the relative number of reads that are aligned to a contig, and chimeras in an assembly make the read mapping more difficult, chimeras alter the accuracy of differential gene expression analysis. For example, if the original sequence of a chimera remains in the assembly, the reads of these transcripts are assigned to the chimera, which likely alters the observed level of expression. In addition, novel transcripts discovery can be complicated by chimeric sequences: 1) chimera can be mistaken for unknown transcripts 2) annotations of new transcripts can be difficult when a contig is composed from multiple transcripts. Removing these chimeras can be a complicated task, because it would require a full transcriptome annotation. This annotation then would have to be screened for genes with double annotations and even then, there is no guarantee that chimeras can be located.

We developed a pipeline that is specifically aimed to filter out chimeras to reduce false-positive gene discovery and false-negative differentially expressed genes. To achieve this goal, we focused on three research aims: 1) to develop a method to assess a *de novo* transcriptome, 2) to use the quality assessment to improve the transcriptome assembly, with specific focus on the removal of chimeric sequences, 3) to make the method as broadly applicable as possible, and to make it as easily applicable as possible. The method of quality assessment and quality improvement is incorporated in an easy to use pipeline with optional user customizability. The pipeline is named after Bellerophon, the hero in Greek mythology that slayed the Chimera. Using Bellerophon to target and remove assembly errors is a useful addition to the short list of transcriptome quality improvement tools currently available.

## Materials and Methods

### The Bellerophon pipeline

To automate the process of filtering and optimizing the transcriptome assembly, the Bellerophon pipeline was developed. This pipeline requires only the sequencing reads and the transcriptome assembly. The user is able to customize the cut-off scores used in the filtration, the order in which to apply the filters and the number of threads used. Bellerophon automatically generates a report that states the results of the pipeline. The user is able to customize the filtering order, but works by default as follows (Figure 1): a) Busco and Transrate are used to establish a ground quality score of the unfiltered assembly. b) Bellerophon filters the assembly using a TPM filter with a default cut-off score of 1. c) TransRate-Q is used again to assess the quality of the new transcriptome. d) Bellerophon then uses CDHIT-EST to filter redundant contigs, using a default cut-off of 95% identity. e) TransRate is executed to use TransRate’s filtering capabilities and to assess the quality of the assembly after TPM and CDHIT-EST filtering. 6) To assess the quality of the fully filtered transcriptome, TransRate-Q and BUSCO are executed using this filtered assembly. Default thresholds were determined using community defaults for TPM (1.0) and ORF length (50 aa or 100aa). For CD-HIT, the default value was determined at 0.95 in order to make the filter more lenient compared to the tool’s default of 0.9. Although these values are set as default by Bellerophon, they can be easily changed to accommodate the user’s dataset.

**Figure 1.**
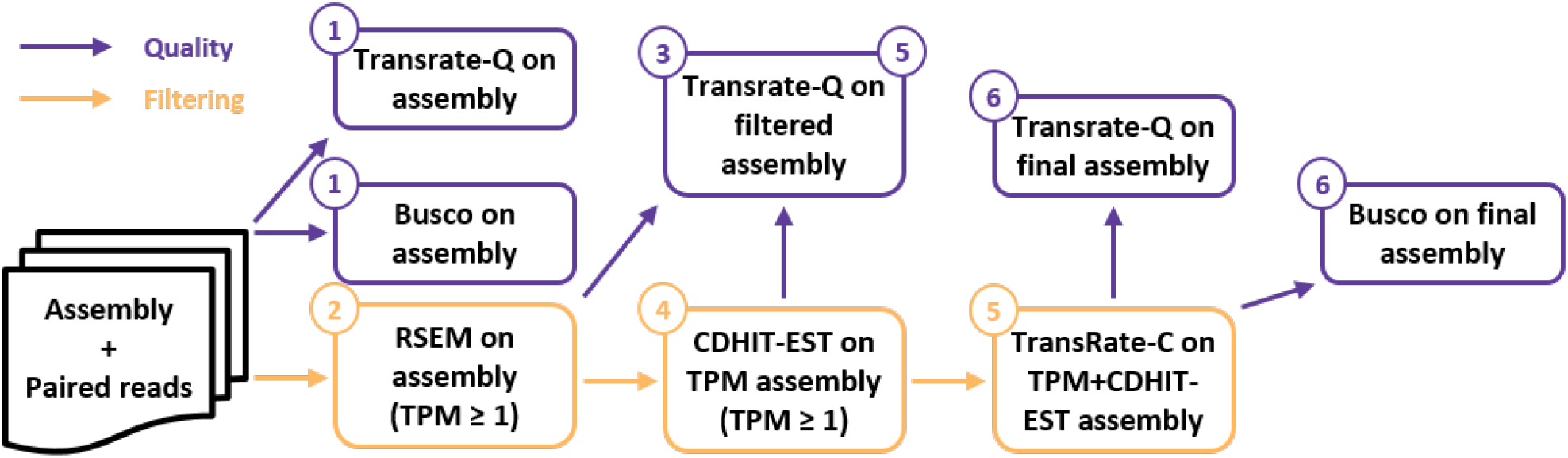
Bellerophon pipeline default filtration order. Violet paths uses TransRate-Q and BUSCO to assess assembly quality. Orange paths displays the sequential filtering steps. The output of each filtering step is used as the input of the following filtering step.

### Validation

To test and validate the Bellepheron pipeline, we used two datasets: 1) RNAseq data obtained by Illumina sequencing the pheromone glands of females from a lab strain of *Heliothis subflexa* (Lepidoptera, Noctuidae), which first needed to be assembled *de novo*, 2) a simulated RNAseq experiment obtained through the tool polyester (Frazee, Jaffe, Langmead, & Leek, 2015) and a reference transcriptome of 3000 transcripts expressed by *Drosophila melanogaster* (Diptera: Drosophilidae).

These datasets where used in three different experiments. The first experiment used the *de novo* assembly of the *H. subflexa* pheromone gland transcriptome and chimeras computationally created from this transcriptome. Details on the assembly of the H. subflexa pheromone gland transcriptome assembly can be found in the supplementary materials. The second experiment focused on the chimeras contained in the ten contigs groups of the *H. subflexa* pheromone gland transcriptome for which the assembler predicted the most isoforms. The third experiment used a Trinity assembly generated from simulated RNAseq experiment of 3000 *D. melanogaster* transcripts. In this experiment, we were able to recognize rightfully assembled contigs from chimera by comparing them with the reference *D. melanogaster* transcripts. Additionally, BUSCO was used to compare the completeness of the *H. subflexa* transcriptome before and after filtering with Bellerophon, using the Insecta odb9 ortholog set.

#### a) Validation using computationally created chimeras

To benchmark the performance of Bellerophon, a set of 500 chimeras was created by randomly selecting two sequences from the *H. subflexa* RNA-seq dataset. These sequences were combined by randomly choosing a percentage between 30 and 70 percent overlap and concatenating the sequences in these proportions. The newly generated chimeras were placed with the other contigs in the assembly. This process was repeated five times. Each assembly with created chimeras was subjected to the Bellerophon pipeline. To test if a significant percentage of chimeras was removed by each step, we compared it to the mean percentage of sequences that was removed by the same step using an unpaired t-test followed by a Bonferroni correction.

#### b) Validation using real assembled chimeras in isoform rich contig groups

Trinity uses an algorithm to find possible isoforms, which occasionally produces more isoforms than actually occur *in vivo*. This makes groups of isoform-rich contig good candidates to search for chimeric sequences. The ten contig groups with the highest number of isoforms were selected and Blasted against the non-redundant protein database (NR), using the BLASTX algorithm (E-value cut-off: 10^−4^). Contigs were identified as chimeric when a sequence contained two matches that did not overlap in their mapping region and had hits with different genes. These contigs are shown to have multiple transcripts on one contig and were marked as a chimera. To measure the ability of the different steps to remove chimeric contigs, we counted the number of marked chimeras left in the assemblies after each steps.

#### c) Validation using a simulated *D. melanogaster* RNA-seq experiment

To further evaluate the performance of the our filtering methods, we used the tool Polyester (Frazee et al., 2015) to generate RNA-seq reads from a random selection of 3000 transcripts from the *D. melanogaster* reference genome (NCBI RefSeq GCF_000001215.4). The expression profile of *D. melanogaster* transcripts was defined through the “fpkm_to_counts” and the “create_read_numbers” functions of polyester, using the expression values of the contigs from our assembly of the *H. subflexa* female pheromone gland as input. 3 sets of 20370192, 19995866 and 20045180 reads were generated using the “simulate_experiment_countmat” function. These reads were assembled using Trinity (Grabherr et al., 2011). Assembled contigs matching less than 5 reads were removed. To determine which assembled contigs were chimeric and which were not, we Blasted the assembled transcriptome against the reference *D. melanogaster* transcriptome (BLASTn, ID percentage cut-off: 90%; e-value cut-off: 10). Contigs matching more than one transcript from the reference transcriptome were considered chimeric. Contigs matching exactly one were not.

#### d) Validation using a *D. melanogaster* RNA-seq experiment

To further evaluate the performance of our filtering methods, we assembled a *de novo* transcriptome using RNA-seq reads, from *D. melanogaster* virgin female heads, available on the Sequence Read Archive of the NCBI (Bioproject: PRJNA527373; experiments: SRR8735410, SRR8735411 and SRR8735412). The transcriptome was assembled following the same protocol as above. We Blasted the contigs assembled by trinity against the reference set of *D. melanogaster* transcripts with the following filters: evalue ≤ 10^−3^, percentage of identify ≥ 90, length of alignment ≥ 300 nucleotides. *D. melanogaster* transcripts were related with their gene of origin. Transcriptome contigs matching only one gene were considered as non-chimeric. Contigs matching multiple genes were considered as chimeric. Contigs matching no genes were considered as unidentified.

## Results

### Bellerophon optimal filtering order

Using the three filter components CD-HIT-EST, TPM and ORF length, an optimal filtering method was designed. To assess the performance of the filter, TransRate-Q was run after each filtering step. Figure 2 displays the TransRate quality score and the number of transcripts remaining after filtering. The filter with the highest resulting TransRate score was the filter with only CD-HIT-EST applied (Figure 2.a, experiment number 3), resulting in a TransRate score of 0.319. However, this filter showed the smallest improvement in the segmented transcripts, leaving 12,096 segmented contigs in the assembly, and a relatively large number of sequences uncovered. The second-best filtering method was the filter using CD-HIT, TPM and ORF-length filters (Figure 2.a, experiment number 6), with a score of 0.316. The filter seemed to improve the most with uncovered and segmented contigs, removing all uncovered contigs and leaving 3,293 contigs in the assembly. However, this filter removed 77,602 (82.9%) contigs, leaving the set with only 16,057 contigs. The third and fourth-best TransRate score were obtained by using CD-HIT-EST first followed by TPM, and TPM first followed by CD-HIT-EST (Figure 2.a, experiments number 10 and 12). These two filtering step removed less contigs overall than the CD-HIT-EST, TPM then ORF-length filtering.

**Figure 2.**
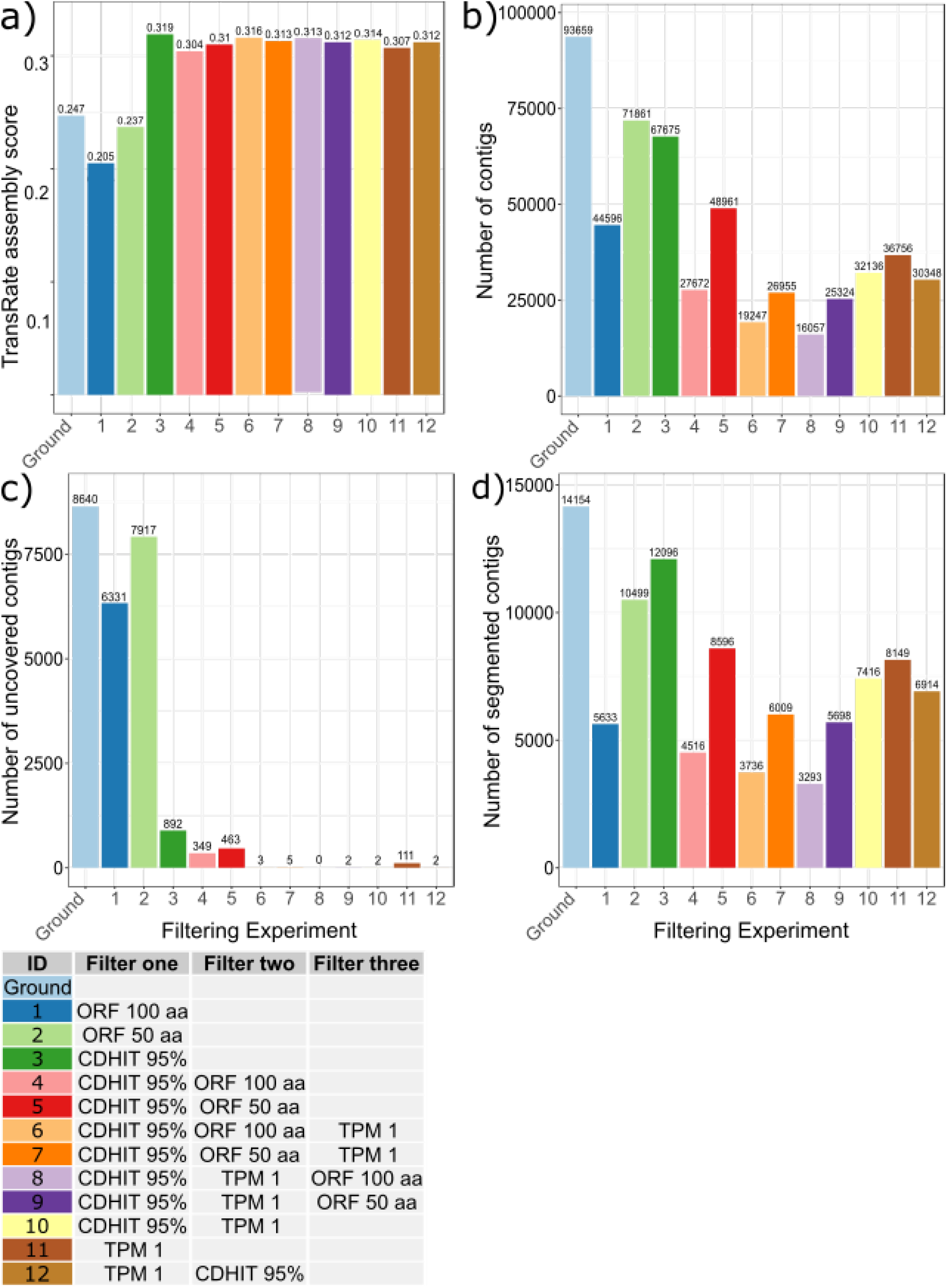
TransRate statistics for every filtering step to find the optimal filtration method. Every letter corresponds to a filtering experiment. The side-table indicates the color and letter coding of the filters applied and their order. Ground bars are made by running Transrate-Q on the unfiltered transcriptome. a) TransRate assembly scores for each filtering experiment. Higher scores indicate higher quality. b) Number of transcripts for each filtering experiment c) Number of transcripts with less than one average per-base coverage per filtering experiment. d) Number of segmented (chimeric) contigs, *i.e*. having un-uniform expression patterns for each filtering experiment.

Since the TPM - CD-HIT-EST filtering reduced the number of segmented contigs in the assembly more than the reverse filtering, we set the default order to be 1) TPM filtering and 2) CD-HIT-EST (filtering order 12 in Figure 2), as the ORF-length filtering appeared to be removing too much transcripts. This was the used order for the chimera removal benchmarking. However, the user of Bellerophon can use any set of filters in any desired order.

### BUSCO and contig length benchmarking

Running BUSCO on the reference transcriptome before and after the Bellerophon pipeline shows a reduction in the number of present groups from 1511 (91.1% of total) to 1420 (85.8% of total). In contrast, the number of duplicated groups is reduced from by 562 to 182 (22.9% of the total number of groups). The BUSCO benchmark numbers are shown in Table 1.

**Table 1.**
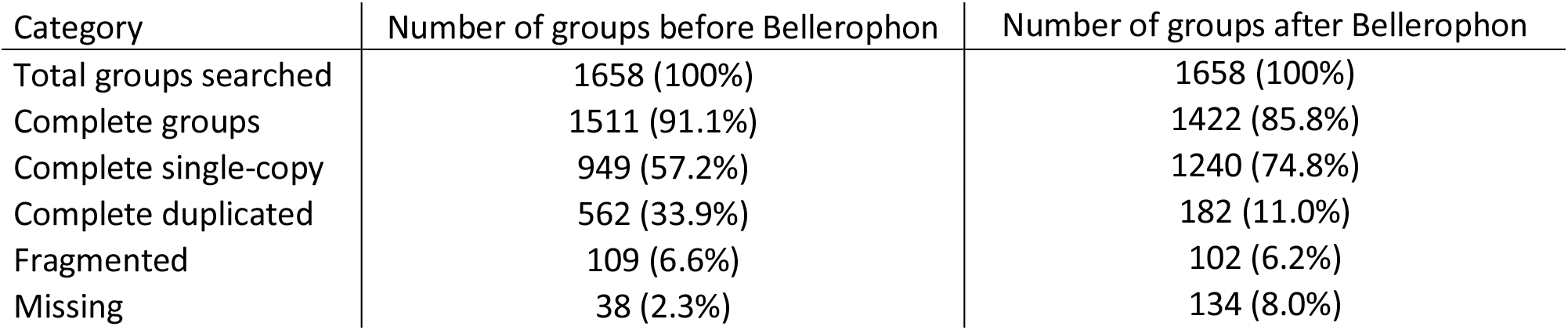
Number of identified BUSCO groups in the *H. subflexa* transcriptome before and after filtering with Bellerophon. Complete single-copy and complete duplicated are subgroups of complete groups. Percentages between parentheses indicate the percentage of the total number of groups.

The length of contigs present in the assemblies or removed from them before and after each filtering step are plotting on the Supplementy Figure S1.a. The mean sequence length is higher for the contigs removed by the TPM and CDHIT filtering test than for the contigs kept by those filters (Table 2). The contigs removed by the final TransRate-C filtering step of Bellerophon are shorter than those that were not removed.

**Table 2.** Mean length of contigs present or removed in the *H. subflexa* and *D. melanogaster* transcriptome by the filtering steps of Bellerophon. The means for *D. melanogaster* are for the *de novo* transcriptome assembled from real RNAseq reads.

|  | Mean contigs length |  |  |  |  |  |
| --- | --- | --- | --- | --- | --- | --- |
|  | *H. subflexa* |  | *D. melanogaster* |  |  |  |
|  | All contigs |  | All contigs |  | Chimera only |  |
|  | Present | Removed | Present | Removed | Present | Removed |
| Pre-Bellerophon | 1385.2 |  | 1474.1 |  | 2267.4 |  |
| Post-TPM | 1358.9 | 1402.2 | 1469.1 | 1685.7 | 2253.9 | 2730.2 |
| Post-CD-HIT | 1329.5 | 1497.9 | 1494.2 | 1352.9 | 2251.3 | 2268.2 |
| Post-Bellerophon | 1369.2 | 817.3 | 1396.8 | 1762.3 | 1910.4 | 2856.4 |

### Validation of filtration of computationally generated chimeras

Out of the total of 500 computationally created and added chimeras to the input assembly, the pipeline removed 485 ± 3.06 chimeras, i.e. 97.04% ± 0.61, of these created chimeras, which was a significantly higher percentage than other sequences that were removed of the input assemblies (69.89% ± 0.03) (unpaired t-test followed by a Bonferroni correction, df = 1, adjusted *P* value = 5.83×10^−6^). Figure 3 shows a flow diagram, displaying the flow of chimeras throughout the experiment. In detail: 1) TPM filtering discarded significantly more chimeras than other sequences (95.72% ± 0.78 vs 60.77% ± 0.00 respectively, *P* = 5.82×10^−6^), 2) the percentage of chimera removed by the CD-HIT-EST filtering was not significantly different from the percentage of other sequences removed (6.94% ± 4.65 vs 17.47% ± 0.01, *P* = 0.35), 3) the Transrate-C filtration step did remove significantly more chimeras than other sequences (26.09% ± 3.63 vs 6.99% ± 0.10, *P* = 0.03).

**Figure 3.**
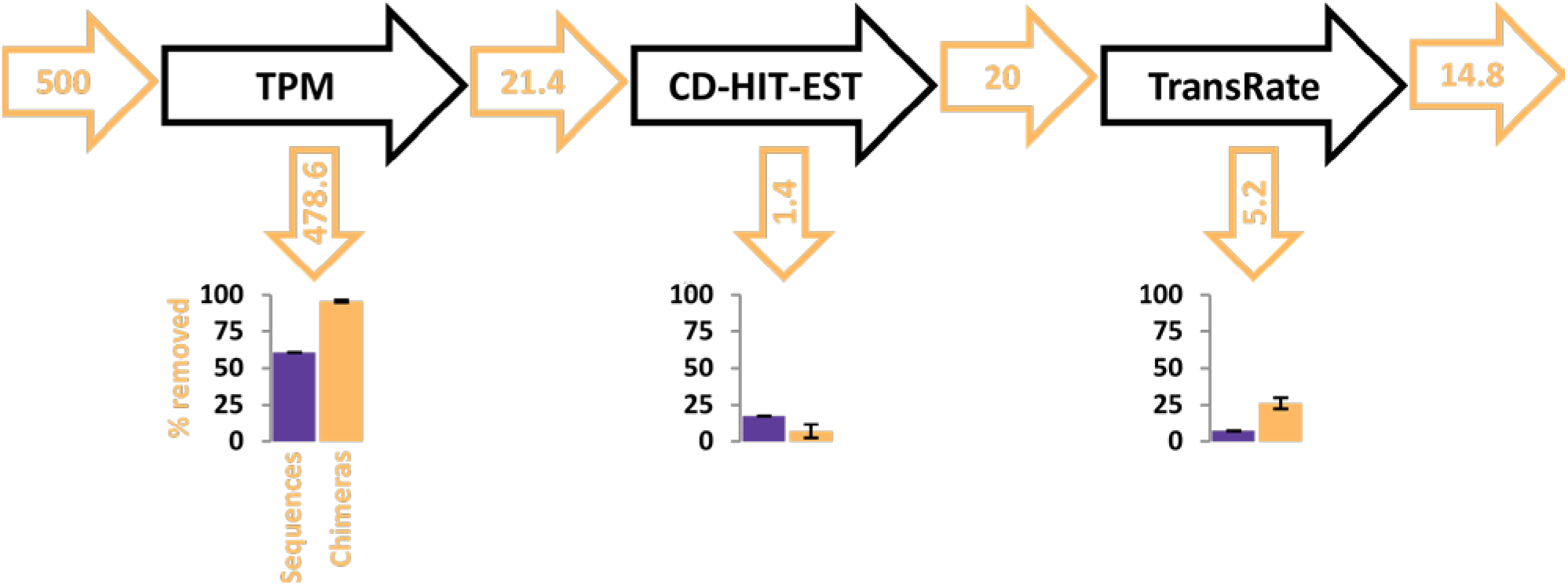
Flow chart of chimeras in the pipeline after testing with 500 intentionally created chimeras. Orange numbers are the means numbers of chimeras in the assembly before and after each filtration steps. Bar charts represent the mean percentage of chimeras and other sequences removed by each step ± SEM.

### Validation using real assembled chimeras in isoform rich contig groups

When focusing on contigs that belong to isoform groups with a large number of contigs, we found that 68 out of 74 (92%) chimeras were removed by the Bellerephon pipeline. Of these 68, 66 were removed by TPM filtering and two by CD-HIT-EST. TransRate-C did not remove any chimera from this set. However, running TransRate-C before running RSEM decreased the number of chimeric isoforms in the benchmark set from 74 to 18, thus removing 56. The quality score of the assemblies in this experiment had increased from 0.247 to 0.336.

### Validation using a simulated *D. melanogaster* RNA-seq experiment

In this simulated RNA-seq experiment, we used 3000 transcripts expressed by *D. melanogaster* as a reference to simulate RNAseq reads. We then assembled those reads with Trinity, which resulted in 3709 contigs. By blasting the Trinity assembly against the reference *D. melanogaster* expressed transcripts, we could relate 3578 contigs to the original 3000 *D. melanogaster* transcripts from which the reads were generated. Through this blasting, we identified 295 contigs of the 3578 contigs as chimeric, and the other 3283 contigs as correctly assembled. Figure 4 shows a flow diagram, displaying the flow of chimeras and other sequences throughout the experiment. The Bellerephon pipeline removed 136 of the 295 chimeras (46.1%), while it removed 575 of the 3283 correct sequences (17.5%). In detail: the TPM filtering step first removed 54 (9.8%) of the chimeras and 349 (8.2%) of the correct sequences; the CD-HIT-EST filtering step removed 30 (12.4%) of the chimeras still present in the assembly at this stage and 64 (2.2%) of the left-over correct sequences. The final Transrate-C filtering step removed 52 (24.6%) chimeras and 162 (5.6%) of the finally left-over correct sequences.

**Figure 4.**
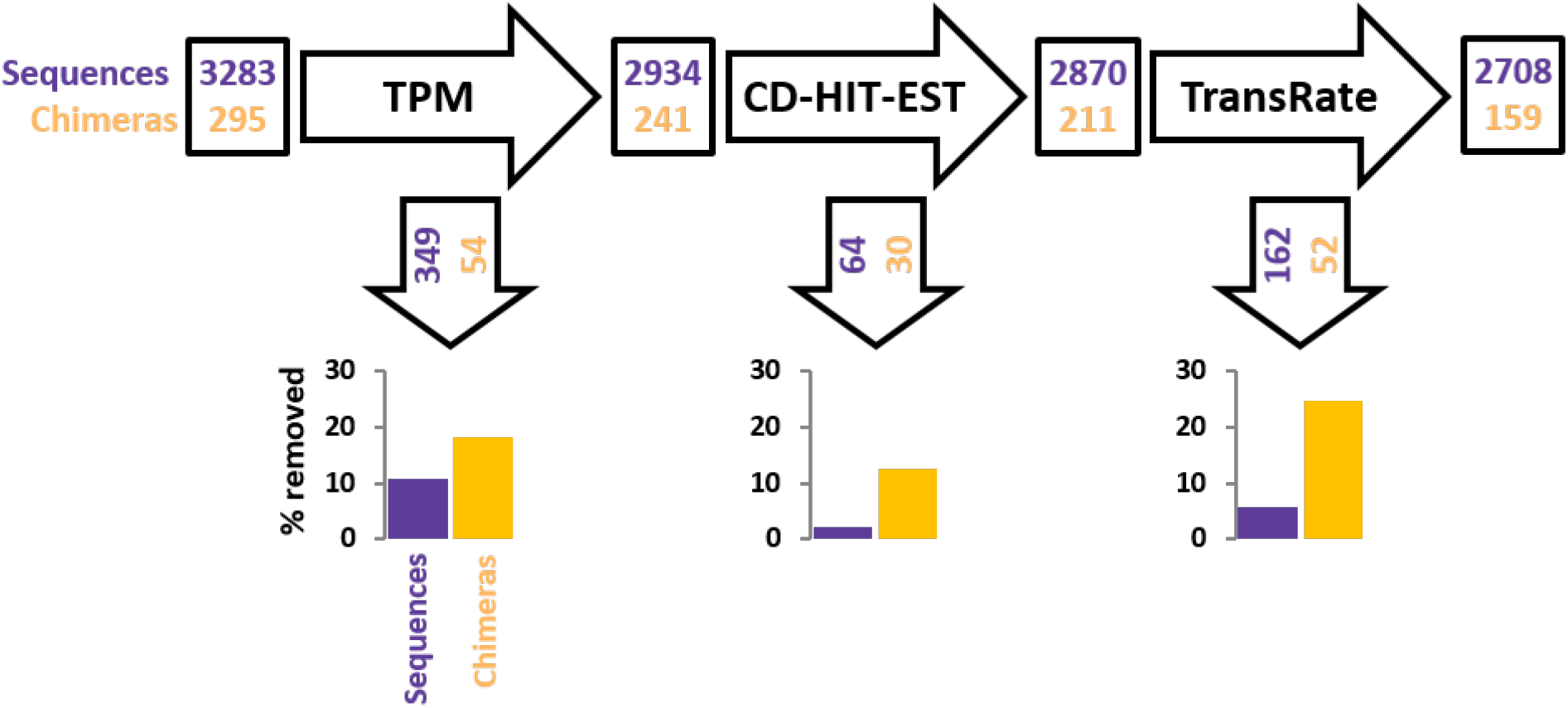
Flow chart of the number of contigs (chimeric or not) during the *D. melanogaster* simulated RNAseq experiment. Numbers in black boxes are the number of chimeras (orange) and other sequences (violet) left in the assembly at the different stages. The numbers in black arrows refer to the number of chimera or other sequences removed by each filtering step. The bar charts display the percentage of chimera (orange) or other sequences (violet) removed by each filtering step.

### Validation using a *D. melanogaster de novo* transcriptome

In this experiment to further test the effect of Bellerophon on *de novo* transcriptome assembly, we assembled a transcriptome of *D. melanogaster* using RNAseq reads available in the sequence read archive. We then assembled those reads with Trinity, which resulted in 33 484 contigs. By blasting the Trinity assembly against the reference *D. melanogaster* expressed transcripts, we could relate 29 201 contigs to *D. melanogaster* transcripts. 26528 contigs matched only on gene and were considered as non-chimeric sequence. 2673 contigs matched multiple genes and were considered as chimera. 5183 contigs didn’t hit any *D. melanogaster* transcripts and remained unidentified.

Figure 5.a shows a flow diagram, displaying the flow of chimeras and other sequences throughout the experiment. The number of correct, chimeric and unidentified contigs in the assembly along the filtering steps are also plotted on the Figure 5.b. The Bellerephon pipeline removed 1619 of the 2673 chimeras (60.6 %) and 3005 of the 5183 (58%) unidentified sequences, while it removed 14325 of the 26528 correct sequences (54%). In detail: the TPM filtering step first removed 227 (8.5%) of the chimeras, 223 (4.3%) of the unidentified sequences and 1952 (7.4%) of the correct sequences; the CD-HIT-EST filtering step removed 600 (24.5%) of the chimeras still present in the assembly at this stage, 1415 (28.5%) of the unidentified sequences and 7057 (28.7%) of the left-over correct sequences. The final Transrate-C filtering step removed 792 (42.9%) chimeras, 1367 (58%) of the unidentified sequences and 5316 (54%) of the finally left-over correct sequences.

**Figure 5.**
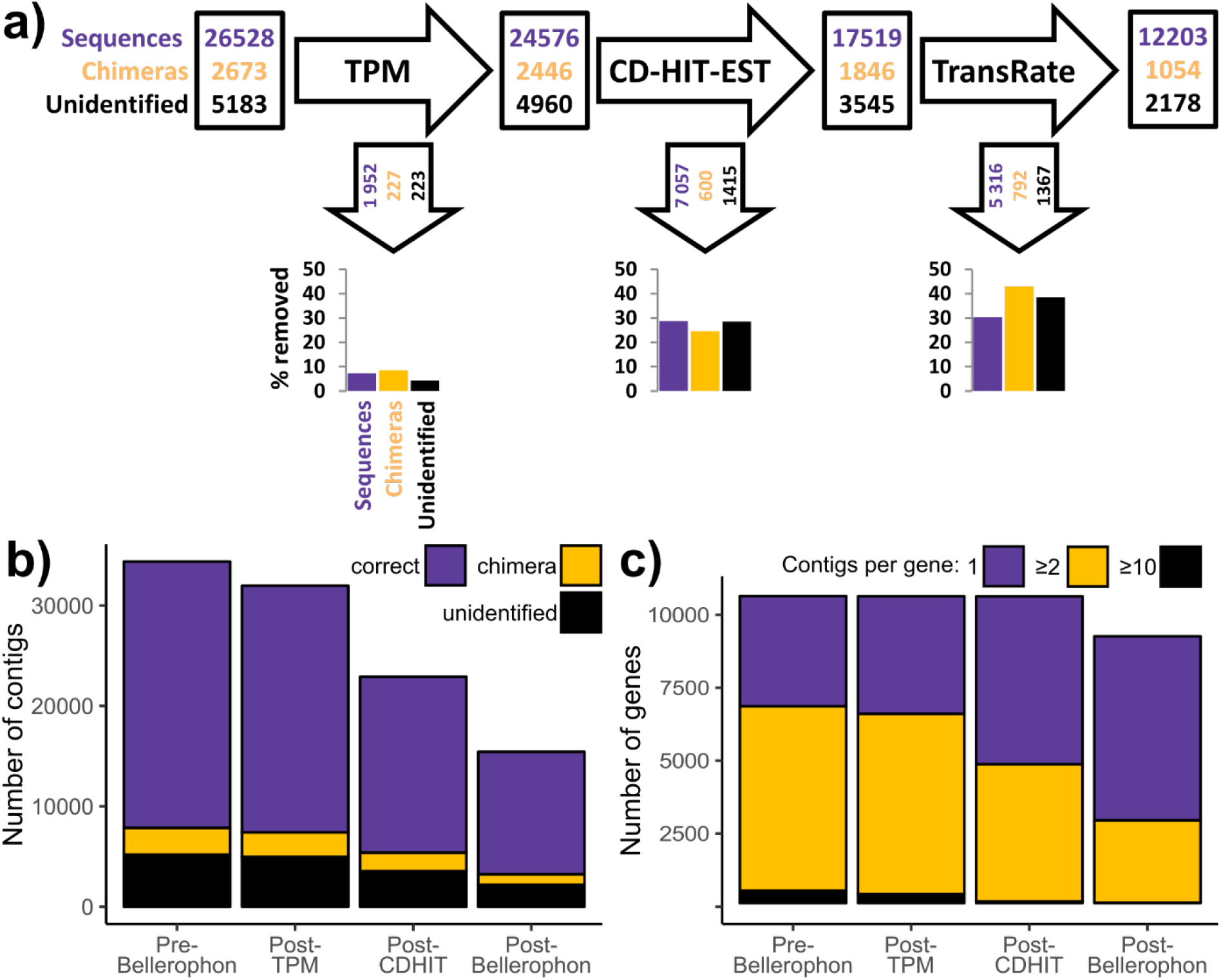
Number of contigs (non-chimeric, chimeric or unidentified) and genes along filtering of the the *D. melanogaster de novo* transcriptome. **a)** Flow chart of the number of chimera, correct and unidentifed sequences filtering by Bellerophone. Numbers in black boxes are the number of sequences present in the assembly at the different stages. Numbers for non-chimeric sequences are displayed in violet, in orange for chimeric (orange) or in black for undidentified sequences (sequences which could not be attributed to a gene through a blast).The numbers in black arrows refer to the number of non-chimeric, chimeric or unidentified sequences removed by each filtering step. The bar charts display the percentage of non-chimeric (violet), chimeric (orange) or unidentified sequences (violet) removed by each filtering step. **b)** Number non-chimeric, chimeric and unidentified contigs along the differents step of the filtering of the *de novo D. melanogaster* transcriptome with the Bellerophon pipeline. **c)** Number of genes represented by at least a contig in the *de novo D. melanogaster* transcriptome with the Bellerophon pipeline. Number of contigs per gene are display in violet (1 contig only), orange (between 2 and 9 contigs) and in black (10 contigs or more).

While the number of contigs matching only one gene in the assembly is reduce by a substantial amount by the Bellerophon filtering, the number of unique genes in the assembly is reduced from 10527 to only 9148 (Figure 5.c). The number of gene represented by only one contig goes from 3778 (35.9% of all gene in the assembly) to 6310 (69%) while the number of genes represented by between two and nine contigs goes from 6327 (60.1%) to 2826 (30.9%).

In order, to check weather Bellerophon removed the shortest sequence of the assemblies, we plotted the number of sequence present and removed at each filtering steps (Supplementary Figure S1.b). The mean length of the contigs kept or removed along the filtering steps of the pipeline are displayed on the Table 2. These values shows that TPM and TransRate-C filtering remove contigs longer than those they keep. Furthermore, the mean length of chimeric contigs are always higher than then mean length of chimeric contigs not filtered out (Table 2).

## Discussion

When there is no reference genome available, the quality of a *de novo* assembled transcriptome is difficult to assess, as there are no indications which transcripts are correctly assembled and which are not. Bellerophon uses the functions of TransRate and three additional post-assembly filtering steps to give insights in the quality of the transcriptome and to maximize this quality. With the selected filtering steps, Bellerophon was able to improve the TransRate quality score of the female *H. subflexa* pheromone gland transcriptome from 0.247 to 0.312. It removed 6,838 contigs without sufficient read evidence in the form of uncovered transcripts, and 7,240 contigs that were considered chimeric as they were not uniformly expressed. Furthermore, Bellerophon removed 83.5% of the benchmark computationally-generated chimeras, 91.9% of the contigs identified as chimera from the isoform-rich contig groups, 46.1% of the chimeras in the simulated *D. melanogaster* RNA-seq experiment and 60.6% of the chimeric contigs of the *D. melanogaster* transcriptome. This proves that Bellerophon improves general transcriptome quality and removes false chimeric sequences. The sequences that Bellerophon removed which were not chimera, present properties that are unwanted for other reasons, such as low read mapping and/or redundancy with other sequences.

Contigs representative of transcripts arising from alternative splicing of one gene might be considered as chimera by Transrate if they are unevenly expressed along their sequence. Bellerophon might thus erroneously remove some of such contigs from the assembly. The most unambiguous way to distinguish chimera from alternative splicing is by comparing the contigs to a reference genome, which is thus problematic when no reference is available. However, filtration of rightfully assembled transcripts arising from alternative splicing should be a minor problem in insects, especially for transcriptomes of one or few tissues, because a) alternative splicing of genes appears to be less common in invertebrates than in vertebrates, with a maximum reported frequencies below 40% (Gibilisco, Zhou, Mahajan, & Bachtrog, 2016; Kim, Magen, & Ast, 2007; E. T. Wang et al., 2008), and b) the majority of alternative splicing events show tissue specificity (Hallegger, Llorian, & Smith, 2010). However, users which have a particular interest in alternatively spliced isoforms should consider not using the CD-HIT-EST filtering step.

When selecting for the filtering order, the RSEM (TPM) filtering step was observed to be the step of the pipeline removing a higher percentage of chimeras than other sequences. This is probably because lowly expressed transcripts have less read evidence, and are thus more prone to assembly errors. Furthermore, chimeric sequences are bound to share read mapping with other contigs of the assembly and as such might appear to be lowly expressed. As this first step removes many chimeras, the performance of the following steps may be reduced because the leftover chimeras may be more difficult to identify. The low number of chimeras discarded by CD-HIT-EST might be explained by the fact that the benchmark-chimeras were randomly selected. CD-HIT-EST works by clustering transcripts based on their sequence identity. The chance of a transcript made up of two randomly chosen transcripts that are 95% identical to another transcript is very low. In the benchmark experiment focusing on assembled chimeras in isoform rich contig groups, the final TransRate-C step of the pipeline did not seem to remove any chimera. The transcripts from these groups presumably belong to one gene family, while a large number of isoforms is created by Trinity. Probably, fewer reads aligned to the false isoforms than to the real isoforms, so that the false isoforms had a low overall expression, increasing the likelihood that the isoforms were removed by RSEM than by TransRate-C runs after RSEM. Our observation that running TransRate-C before running RSEM decreased the number of chimeric isoforms in the benchmark set from 74 to 18, removing 56 chimeras, confirms this suggestion.

A good comparison between different available tools, i.e. KisSplice, DRAP, RSEM-EVAL, TransRate and Bellerophon, is difficult, because there are great differences in used datasets between the different studies. Overall, the number of chimera that we found is much higher than those found in other studies. Bellerophon found 5,053 (17.9%) chimeras in its final assembly. In comparison, Lima *et al* labelled 1.3 % of their contigs created with KisSplice as chimeric (Lima et al., 2017), similar to Cabau *et al.* who found 0.09 % to 0.56 % chimeras in Trinity assemblies (Cabau et al., 2017), while Yang and Smith found approximately 4 % chimeric sequences among Trinity assembled contigs (Yang & Smith, 2013). All studies have used a different way to pinpoint chimera: KisSplice uses an algorithm that is based on the percentage of mapped reads that match, while Cabau *et al.* and Yang & Smith used a self-alignment method in transcripts to find chimera, using simulated data based on well-referenced datasets of *Homo sapiens* and *Danio rero*. The assembly used in our research contained 93,659 contigs, and the reads were only from one tissue: the pheromone gland of the moth *H. subflexa*. As the full transcriptome of the moth model *Bombyx mori* contained 37,408 transcripts (Li et al., 2012), the high number of transcripts in our dataset shows an over-prediction of isoforms and other assembly errors by Trinity. The *D. melanogaster* transcriptome filtering allowed us to observe that gene representation is only marginally impact by Bellerophon. Eliminating chimeras and other assembly errors has made our dataset cleaner and more optimized for further differential expression analysis of RNA-seq experiments.

## Acknowledgements

We thank David G. Heckel for his comments on the manuscripts, Rik Lievers for his help in preparing moth tissue samples and extracting RNA, and the BIPAA bioinformatics platform (Rennes, France) for hosting some of the computations conducted. This project was funded by INRA, the Netherlands Organisation for Scientific Research (NWO-ALW, award no. 822.01.012) and the National Science Foundation (award no. IOS-1052238 and IOS-1456973).

## Data Accessibility Statement

The raw reads used to assemble the *Heliothis subflexa* pheromone gland transcriptome available in GenBank SRA under the ID: PRJNA493752.

The *D. melanogaster* transcript used for the simulated RNA-seq experiment were downloaded from GenBank genome assembly number <u>GCF 000001215.4</u>.

The reads used to assembled the *de novo D. melanogaster* transcriptome are available in SRA under the IDs: SRR8735410, SRR8735411 and SRR8735412

The Bellerophon pipeline in available at Github at the link: https://github.com/JesseKerkvliet/Bellerophon.

## Author Contributions

JK, AdF, MvW and AT designed the research.. JK and AdF performed the research. All authors wrote the paper.

