## Supplementary Materials for "The Bellerophon pipeline, improving de novo transcriptomes and removing chimeras"

### Supplementary Material and Methods

#### Transcriptome of the pheromone gland of the noctuid moth *Heliothis subflexa*

##### a. Sequencing and assembly of *H. subflexa* pheromone gland transcriptome

Using the popular *de novo* transcriptome assembling tool Trinity (Haas et al., 2013), we assembled the RNA-seq reads obtained by Illumina sequencing of 10 female *H. subflexa* pheromone glands. After assembly, we removed all contigs from the assembly that did not align with 5 or more reads. This reference assembly is used as reference transcriptome for the quality assessment and chimera filtration. To have an indication of the completeness of this assembly, the Benchmarking Universal Single-Copy Orthologs (BUSCO) tool was used (Simão, Waterhouse, Ioannidis, Kriventseva, & Zdobnov, 2015). This tool searches for a reference set of genes that should be present in every insect and should occur only once.

##### b. Transcriptome quality assessment

To assess the quality of a *de novo* assembled transcriptome, we used the tools TransRate (Smith-Unna et al., 2016) and BUSCO (Simão et al., 2015). TransRate uses the assembly and the raw paired-end reads that were used to make the assembly. Based on the mapping data, it calculates four different quality scores that, when combined, indicate the quality of the assembly. This combined quality score is a figure between zero and one. The higher this score, the higher the quality. When using TransRate as a quality measuring tool (TransRate Score), it is referred to as Transrate-Q. Busco is a tool that search for conserved single-copy genes in the transcriptome input. Busco report if those genes are found complete or partial and in single copies or duplicated copies.

##### c. Transcriptome filtering methods

To reduce the number of chimeric contigs in the transcriptome assembly, we used different methods and tools. First, the tool RSEM (Li & Dewey, 2011) was used to calculate the relative expression of contigs, using the Transcripts Per Million (TPM) metric, and to remove lowly expressed contigs. These contigs are thus contigs for which the assembler did not have much read evidence. The used TPM cut-off was 1.0. The second tool used was CD-HIT-EST (Fu, Niu, Zhu, Wu, & Li, 2012) that clusters sequences based on similarity. For every cluster, a representative sequence is chosen, while the others are discarded. This filter step thus removes redundancy from the assembly. The identity cut-off was 95%. Third, TransDecoder, a part of Trinity, was used to remove contigs containing ORFs of fewer amino-acids than the set cut-off score. Cut-off scores were set at 50 aa and 100 aa. Finally, we use TransRate as a filtering tool, therefore referred as TransRate-C (for Chimera). Is this case, TransRate relies on quality scores it measures to filter contigs from the input assembly. To find the optimal combination of filters, different filter orderings were applied to the assembly.

### Supplementary Results

#### Transcriptome assembly quality

Sequencing the *H. subflexa* cDNA resulted in a total yield of 120,107,603 paired-end reads per individual. The per-sample yield is displayed in Supplementary Table S1. Assembling with Trinity resulted in a total of 93,659 contigs, deriving from 35,500 putative genes. Supplementary Table S2 displays the average transcript length, median transcript length, N50, GC percentage and total number of bases. An initial TransRate run gave a base Assembly Score of 0.2465. Using BUSCO, a set of single-copy orthologues was searched that every insect should contain. This resulted in a set of 1,511 orthologues found, out of a total 1,658 orthologues defined by BUSCO (i.e. 93%). Of these, 949 (63%) were found as single-copy. The results of BUSCO are displayed in Table 1.

#### Supplementary Figures

**Supplentementary Figure S1. Repartition of contig length of filtered and unfiltered sequences during filtering of the *H. subflexa* and the *D. melanogaster* transcriptomes.**

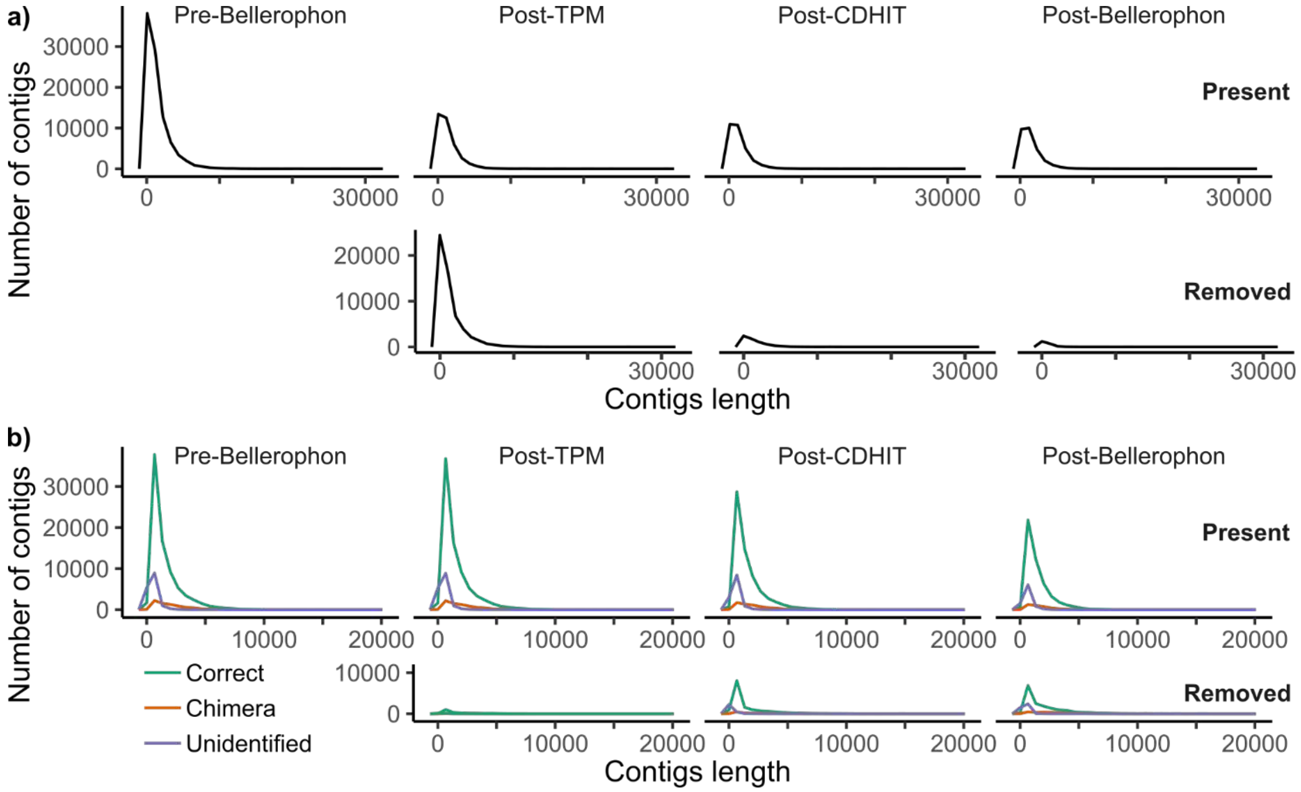


Repartition of contig length that are present or have been removed before filtering and after TPM, CDHIT and TransRate-C filtering for: **a**) *H. subflexa de novo* transcriptome and **b**) *D. melanogaster de novo* transcriptome assembled from real RNAseq reads. The contig length of correctly assembled, chimeric and unidentified contig is display, respectively, in green, orange and violet for the *D. melanogaster* transcriptome.

#### Supplementary Tables

**Supplementary Table S1. Number of pair end reads for each sample.**

Number of paired-end reads couple in each RNAseq samples used to assemble the *Heliothis subflexa* pheromone gland transcriptome.

| Sample number | Number of paired-end reads |
|---|---|
| 14 | 10,709,318 |
| 36 | 10,285,139 |
| 39 | 17,653,014 |
| 48 | 9,727,593 |
| 87 | 14,795,805 |
| 22 | 10,440,250 |
| 35 | 10,988,040 |
| 38 | 10,663,367 |
| 53 | 15,417,091 |
| 88 | 9,427,986 |

**Supplementary Table S2. Assembly statistics for the *Heliothis subflexa* assembly.**
Various metrics and statistics of the *Heliothis subflexa* pheromone gland transcriptome assembly.

| Statistic | Number |
|---|---|
| Number of contigs | 93659 |
| Number of contigs with ORF | 355 |
| Average length of transcripts | 1385.21 |
| N50 | 2571 |
| GC% | 40% |
| Number of bases | 129,737,226 |
| TransRate Score | 0.2456 |
